# Jagged-1 regulates *Foxp3* expression and cytokine production in CD4^+^ T cells

**DOI:** 10.1101/712430

**Authors:** Soichiro Kimura, Ronald Allen, Melissa Scola, Nicholas W Lukacs, Steven L. Kunkel, Matthew Schaller

## Abstract

Notch ligands are present during the interactions between T cells and dendritic cells (DC) and induce a myriad of effects that facilitate the activation of T cells, including the induction of T cell regulation, survival, and cytokine production. Although the ligands Delta-like 4 and Delta-like 1 are expressed as a function of DC activation, the notch ligand Jagged-1 is constitutively expressed on DC. We sought to determine the role of Jagged-1 in the interactions between CD4^+^ T cells and DC. We observed that Jagged-1 regulates Foxp3 expression, and *Cd11c*^*Cre+*^*Jagged*^*ff*^ mice have an altered expression of Foxp3 in effector cells that arise as a result of infection with the mycobacterium *Bacille Calmette-Guerin*. The observed changes in *Foxp3* expression were correlated with an increase in cytokine production from cultures of antigen-stimulated draining lymph nodes.

## Introduction

The Notch system consists of 4 receptors and 5 ligands that interact to determine cell fate and cause cellular differentiation. Studies have suggested that the Delta and Serrate/Jagged-1 families may send different signals through the same receptor^1^. Differential signaling has been observed at the level of gene expression, cell morphology^2^ and cell phenotype^3,4^. Within the immune system, Notch is used by dendritic cells (DC)^5^ and structural cells^6^ to facilitate the activation of T cells during the process of antigen presentation. Although extensive characterization of the role of DLL4 has been performed in regards to T cell differentiation^7– 9,10,11^, less is known concerning the role of the Notch ligand Jagged-1 in this same system. Jagged-1 is constitutively expressed on the surface of DC, whereas DLL4 is induced by MyD88 through TLR stimulation^12^. As a ligand that is expressed on DC at times when no inflammation is present, it is possible that this ligand may cause an immunosuppressive effect. This hypothesis is upheld by several studies that have suggested that Jagged-1 causes the induction of a CD4^+^CD25^+^Foxp3^+^ T-regulatory (Treg) cells^13,14^.

To characterize the contribution of Jagged-1 to T cell activation and differentiation, we used a combination of *in vitro* and *in vivo* approaches. We observed that recombinant Jagged-1 down-regulated the expression of Notch target genes and the transcription factor Foxp3 under conditions that caused the differentiation of T-regulatory cells (iTregs). In other studies, we observed that co-culture of naïve CD4^+^ T cells with DC from *Cd11c*^*Cre+*^*Jagged1*^*ff*^ mice increased the number of Foxp3^+^ cells, thus confirming our data using recombinant protein. *In vivo* we observed an increase in T regulatory cell development during intratracheal infection with the *Bacille Calmette-Guerin* (BCG) strain of mycobacteria at 10 weeks post infection. This increase in Treg numbers was accompanied by an increase in cytokine production when cells from the draining lymph node were stimulated *in vitro* with cognate antigen. These data suggest that dendritic-cell-derived Jagged-1 may regulate *Foxp3* expression and Treg-mediated suppression of immune responses during BCG infection.

## Results

### Jagged-1 decreases Foxp3 expression in CD4^+^ T cells

Previous studies have demonstrated that DLL4 can increase and stabilize *Foxp3* expression in naïve CD4^+^ T cells^9^. To determine the effect of Jagged-1 on CD4^+^ T cell expression of Foxp3, we stimulated naïve T cells with equimolar concentrations of recombinant DLL4 and recombinant Jagged-1. We observed that neither ligand was able to upregulate *Foxp3* expression in T cells in the absence of TGF-ß1 (Th0 condition). We also observed that DLL4 caused no significant change in *Foxp3* expression at the 10 nM concentration under T-regulatory cell conditions (TGF-ß1 + IL-2). Conversely, 10 nM Jagged-1 decreased *Foxp3* expression under these same conditions (fig 1a). We also observed a similar pattern of expression in other Notch target genes, including *Tcf7, IL2RA* (CD25), and *Notch2* (fig S1). We next tested if rJagged-1 decreased *Foxp3* expression in a dose-dependent manner. Stimulation of naïve T cells isolated from *Foxp3*^*eGFP*^ mice under iTreg conditions demonstrated that the reduced expression of *Foxp3* was dependent on the dose of Jagged-1 (fig 1b). Higher doses of rJagged-1 (50 nM) caused a decrease in viable CD4^+^ T cells in the culture (S1). To determine if the decrease in Foxp3 expression was linked with changes in CD25 expression, we assessed CD25 by flow cytometry. Consistent with the PCR data in figure S1, we observed dose-dependent decrease in the number of CD25^+^CD4^+^ T cells in iTreg cultures treated with increasing doses of rJagged-1 (fig 1c). We also observed that stimulation of T cells under iTreg conditions caused a significant decrease in *Foxp3* expression at both 24 and 48 hours post stimulation in those cells treated with 10 nM rJagged-1 (fig 1d).

**Figure 1:**
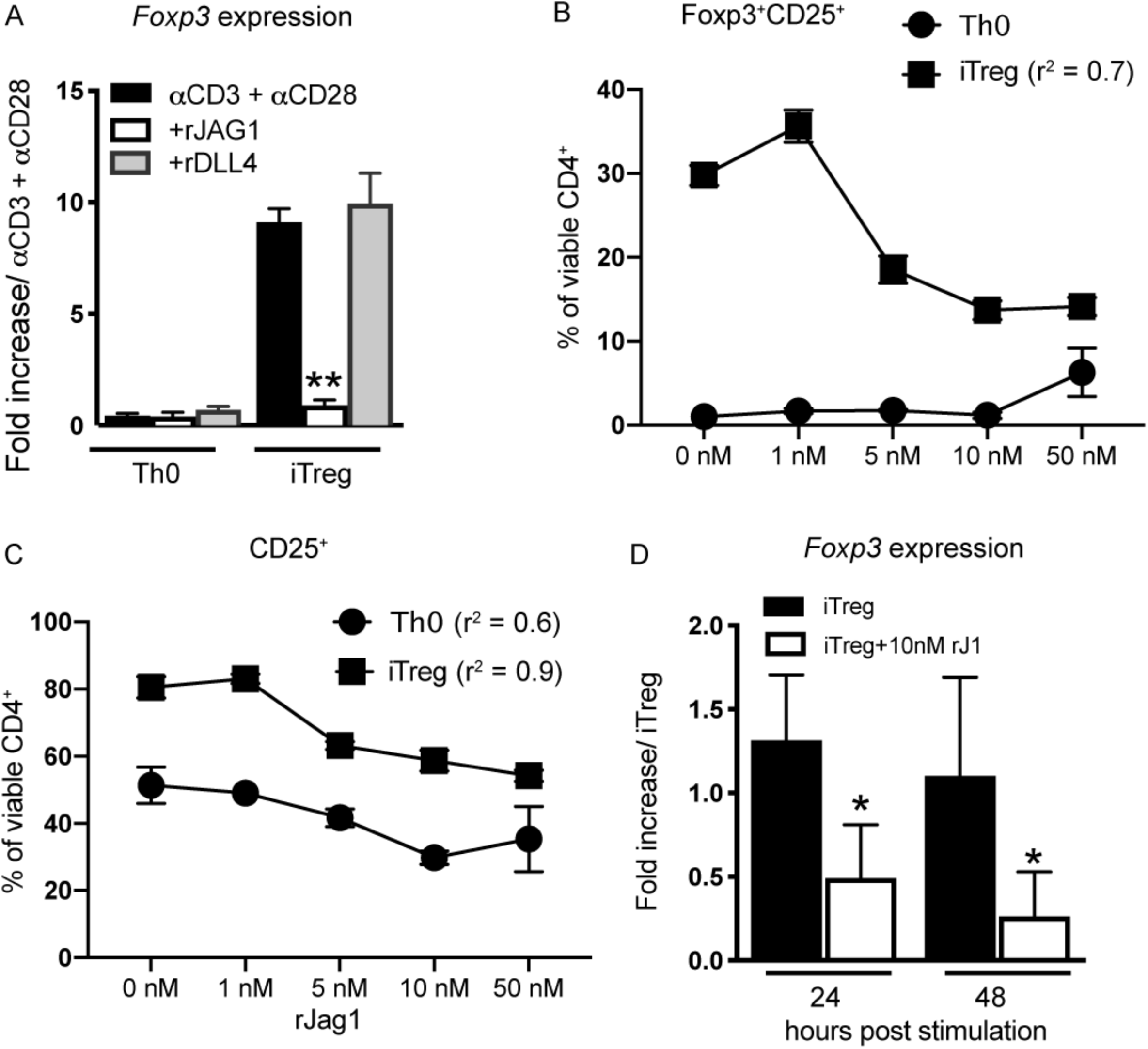
Jagged-1 regulates Foxp3 expression in CD4^+^ T cells. A) Naïve CD4^+^ T cells were stimulated with plate-bound anti-CD3 and anti-CD28 alone (Th0) or in the presence of 10 ng/mL recombinant IL-2 and 2 ng/mL recombinant human TGF-β (iTreg) with or without 10 nM plate-bound rJ1 or rDLL4. *Foxp3* expression was measured at 72 hours post stimulation. B) Naïve CD4^+^ T cells from *Foxp3*^*eGFP*^ mice were isolated and stimulated under Th0 or iTreg conditions with varying concentrations of recombinant rJ1. Flow cytometry was performed at 72 hours post stimulation for analysis of Foxp3^+^CD25^+^CD4^+^ cells. R values were determined using nonlinear regression. C) The percentage of viable CD4^+^ cells that are CD25^+^ cells in the same conditions as outlined in 1B. R values were determined using nonlinear regression. D) Naïve CD4^+^ T cells from C57/BL6 mice were stimulated under iTreg conditions with 10 nM plate-bound rJ1 for varying amounts of time. A two-way ANOVA was used to determine significance, with an overall p-value of 0.0003. In figure 1D, all of the significance was derived from the treatment groups, and none from the time variable.

### Dendritic cells lacking Jagged-1 increase Foxp3 expression in naïve T cells

To determine if DC-derived Jagged-1 would cause the same phenotype as the recombinant Jagged-1 used in our initial experiments, we bred a *Jagged*^*ff*^ strain with *Cd11c*^*Cre*^ to create *CD11c*^*Cre+*^*Jagged*^*ff*^ mice. We FACS sorted CD11c^+^ expressing myeloid cells from the lungs of these mice to test for a reduction of *Jagged1* expression by qPCR. We observed that recruited monocytes and CD103^+^ DC in the lungs of *CD11c*^*Cre+*^*Jagged*^*ff*^ mice had greater than 50% reduction in *Jagged1* expression. Conversely, CD11c^+^CD11b^+^ DC and alveolar macrophages isolated from the lungs of *Cd11c*^*Cre+*^*Jagged*^*ff*^ mice had less than 10% reduction in *Jagged1* expression compared to *Cre*^−^ littermate controls (fig 2a). Splenic DC isolated from *Cre*^*+*^ mice showed a reduced production of the cytokines IL-6 and the IL-12 p40 subunit when stimulated with CpG, but not other TLR stimuli (Fig S2), suggesting these DC were functionally normal in their response to PAMPs. There was no change in DC numbers in the lung, spleen or thymus in naïve mice.

**Figure 2:**
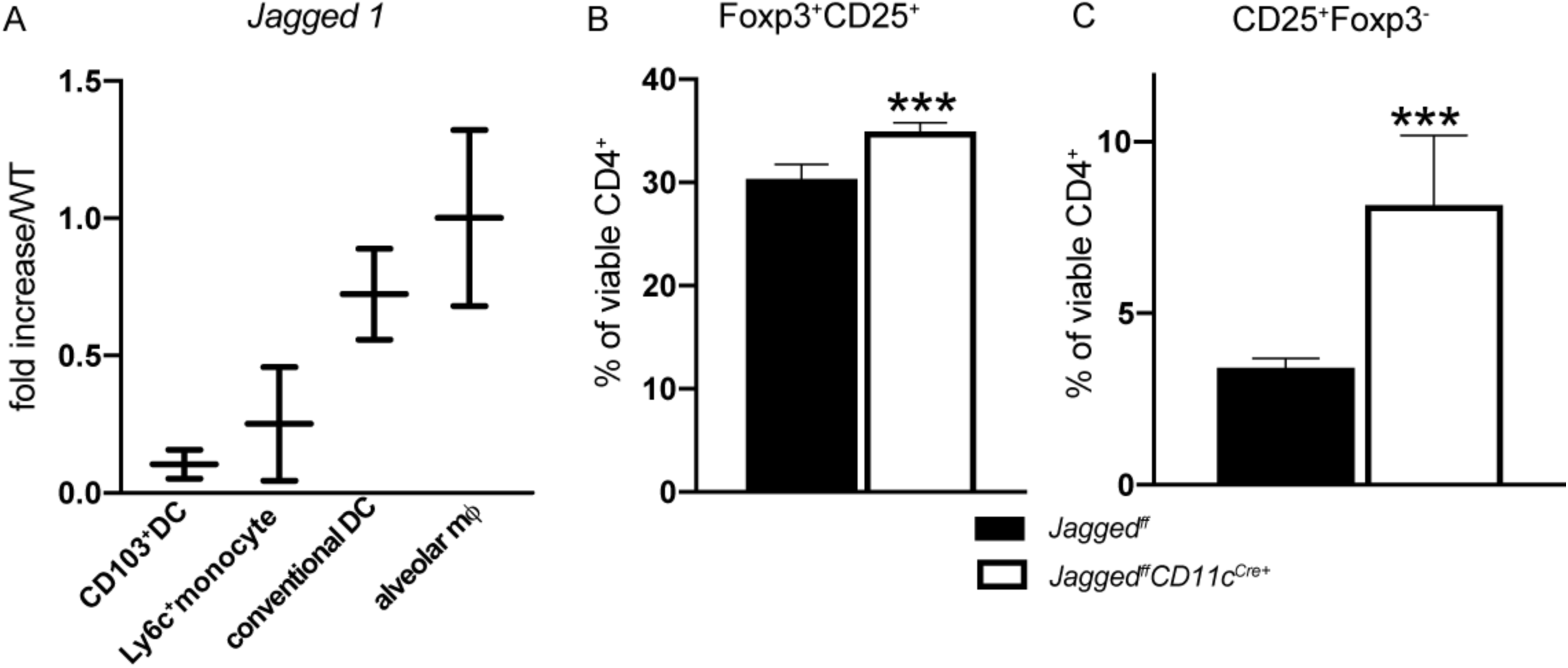
DCs from Jagged1^ff^Cd11c^Cre+^ mice increase Foxp3 and CD25 expression in naïve T cells. A) DC and macrophage populations were FACS sorted from the lungs of *Cd11c*^*Cre+*^*Jagged1*^*ff*^ mice or *Jagged1*^*ff*^ controls and assessed for expression of *Jagged1* by QPCR. Results are depicted as expression relative to the levels measured in *Jagged1*^*ff*^ littermate controls. CD103^+^ DC were gated as autofluorescent^−^CD11b^−^CD11c^hi^CD103^+^, monocytes were autofluorescent^−^CD11c^lo^SSC^lo^CD11b^+^Ly6C^+^, cDC were autofluorescent^−^ CD11c^hi^Ly6C^hi^. B-C) Splenic CD11c^+^CD11b^+^ DC were FACS sorted and cultured with FACS sorted CD62L^+^CD44^−^CD25^lo^Foxp3^−^ naïve T cells isolated from *Foxp3*^*eGFP*^ mice for 7 days in the presence of 3μg/mL soluble anti-CD3. Foxp3 expression was assessed by GFP expression at 7 days after culture. Results were determined to be significant by Student’s t-test (p=0.001 for B and p=0.002 for C).

We next determined if reduced expression of *Jagged1* on DCs could upregulate Foxp3 in iTreg cultures. To do this, we co-cultured naïve T cells isolated from *Foxp3*^*eGFP*^ mice with FACS-sorted splenic Cd11c^+^Cd11b^+^ DC from *Cd11c*^*Cre+*^*Jagged*^*ff*^ mice in the presence of soluble CD3. Co-culture of naïve T cells with DCs isolated from *Jagged*^*ff*^*CD11c*^*Cre+*^ mice caused a significant increase in *Foxp3* expression in CD4^+^ T cells (fig 2b). We also observed a significant increase in the number of CD25^+^Foxp3^−^ cells in co-cultures containing DCs from *CD11c*^*Cre+*^*Jagged*^*ff*^ mice (fig 2c). Analysis of cell culture supernatants from these cultures indicated no significant changes in IFN*γ*, IL-10 or IL-12 (fig S2).

### Increased expression of Foxp3 in Cd11c^Cre+^Jagged1^ff^ infected with BCG

We reasoned that DC:T cell interactions would be relevant during an infectious model, which facilitates the activation of T cells. We chose to infect mice with BCG as signaling via the Notch pathway is an important component of T cell activation in this model^15,16^, and T cells contribute the to the BCG immune response^17^. To examine *Foxp3* expression in *Cd11c*^*Cre+*^*Jagged*^*ff*^ mice, they were crossed with a *Foxp3*^*eGFP*^ strain to generate *Cd11c*^*Cre+*^*Jagged*^*ff*^*Foxp3*^*eGFP*^ mice.

We observed no change in the percent of CD4^+^Foxp3^+^CD25^+^ cells in the lungs or draining lymph nodes at 2 or 5 weeks post infection. However, we did observe an increase in the Foxp3^+^CD25^+^ population at 10 weeks post infection in both the lung and lymph node (fig 3a-b). We observed no change in bacterial growth in *Cd11c*^*Cre+*^*Jagged1*^*ff*^ mice compared to *Jagged1*^*ff*^ controls at any time point (fig 3c), and all mice survived the infection up to 10 weeks after inoculation. To determine if the observed increase in CD4^+^Foxp3^+^CD25^+^ cells was accompanied by altered cytokine production, we stimulated lymph node cells from *Cd11c*^*Cre+*^*Jagged1*^*ff*^ or controls with purified protein derivative (PPD) after 10 weeks of infection. We found that cell culture supernatants from *Cd11c*^*Cre+*^*Jagged1*^*ff*^ contained more IFN*γ* and IL-10 than *Jagged1*^*ff*^ controls (fig 3d).

**Figure 3:**
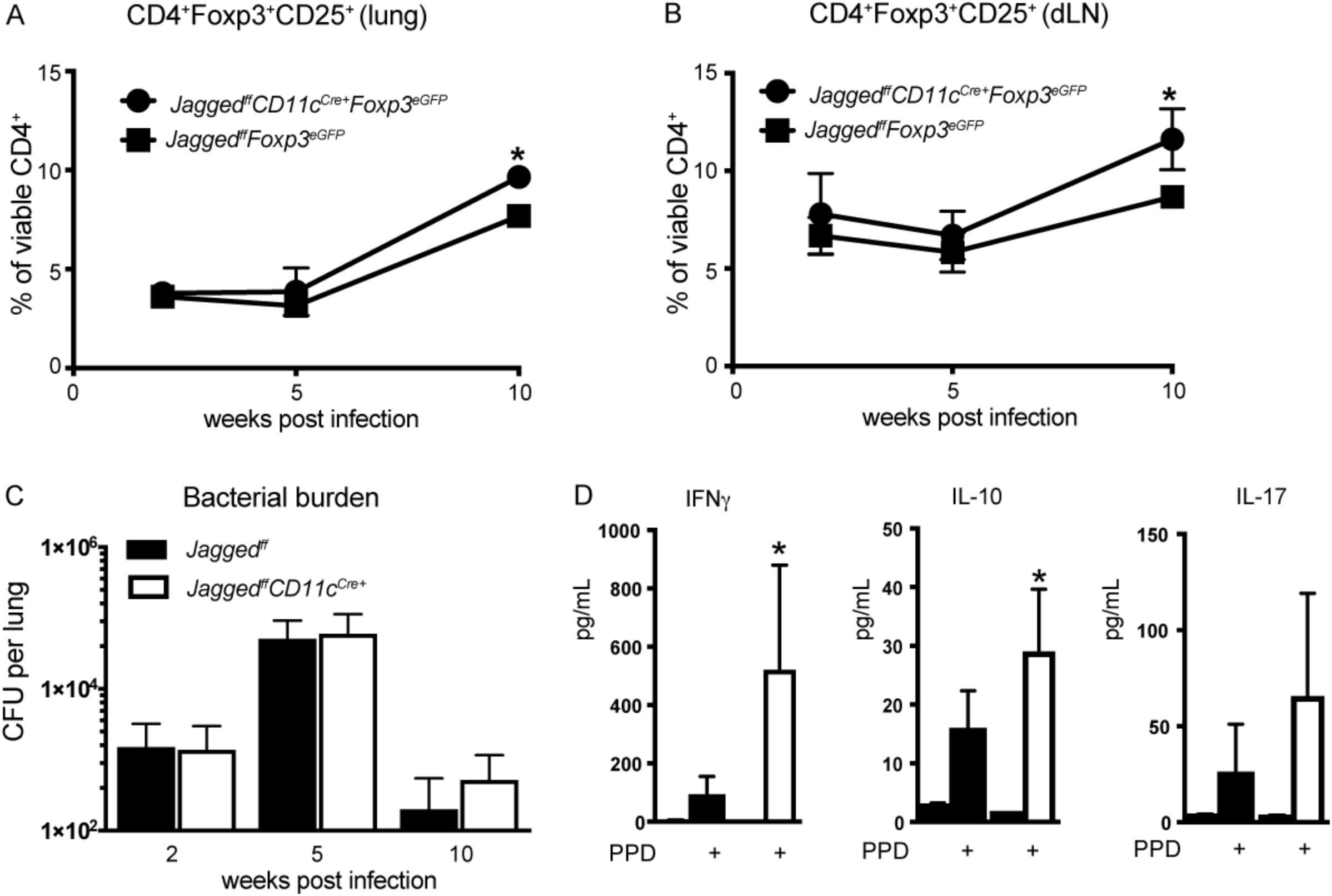
T cell populations and cytokine production altered in BCG infected Cd11c^Cre+^Jagged1^ff^ mice. A-B) Analysis of viable CD4^+^Foxp3^+^CD25^+^ cells in the lungs and draining lymph nodes of *Jagged*^*ff*^*Foxp3*^*eGFP*^ and *Jagged*^*ff*^*CD11c*^*Cre+*^*Foxp3*^*eGFP*^ mice at 2,5, and 10 weeks post BCG infection. *p=0.0007 for lung and *p=0.009 for lymph node as determined by two-way ANOVA. Each time point contained 4-8 mice. C) Colony forming units were measured in the lungs of *Jagged1*^*ff*^*CD11c*^*Cre+*^ mice and littermate *Jagged*^*ff*^ controls at 2, 5, and 10 weeks post infection. D) Cytokine levels from lung-draining lymph node cells from mice at 10 weeks post infection. Single-cell suspensions were stimulated with 10 ug/mL of purified protein derivative (PPD) after 72 hours of *in vitro* culture. Significance was determined using one-way ANOVA analysis. *p-adjusted=0.002 for IFNγ and *p-adjusted=0.005 for IL-10. IL-17 levels were not significantly different between groups.

## Conclusions

The Notch ligand Jagged-1 is generally associated with a suppression of T-cell mediated immune responses, including the generation of T-regulatory cells^14^ and inhibition of Th17 differentiation^18^. A limitation of these previous studies is that they were conducted *in vitro*. We sought to bridge the gap between *in vitro* and *in vivo* studies with the generation of a mouse that specifically eliminated *Jagged1* expression on dendritic cells. Our data indicated that the expression of the *Jagged1* gene was reduced to varying degrees in different populations of CD11c^+^ cells.

Our *in vitro* data suggested that Jagged1 reduced the expression of *Foxp3* at 10 nM concentration, while Delta-like 4 had no effect at this same concentration. Previous studies have indicated that Dll4 can increase Foxp3 expression^9,19^. These previous studies were performed using a 50 nM concentration of Dll4. There are many factors that govern the affinity of Notch ligands for their receptors, including glycosylation^20^ and the expression of Notch receptors, which can change in leukocytes as cells differentiate^21,22^. It is likely these factors influence the affinity and strength-of-signal for Notch ligands as they contribute to Treg differentiation.

We also found that the co-culture of CD4^+^ T cells with DCs lacking *Jagged1* increased the expression of *Foxp3* under iTreg conditions. These data were supported by *in vivo* results demonstrating an increase in Foxp3^+^CD25^+^ in the lung and lymph node of *Cd11c*^*Cre+*^*Jagged*^*ff*^*Foxp3*^*eGFP*^ at 10 weeks post BCG infection. At the same time point that we observed an increase in production of IL-10 and IFN*γ* from antigen-stimulated draining lymph nodes. These data suggest that Jagged-1 may not only down-regulate Foxp3 expression, but also lend stability to the Treg phenotype. Destabilization of Tregs coupled with increased expression of IFN*γ* in the presence of Foxp3 has been observed in other systems including *Helios*^*-/-*^ Tregs^23,24^, *Eos*^*-/-*^ Tregs^25^, and disruption of other intracellular signaling pathways^26^. It is also possible that Jagged-1 down-regulates the Foxp3 expression associated with T cell differentiation from naïve to effector or memory T cells^27,28^. Variations in Jagged-1 expression have been reported in human monocyte-derived DC and inversely correlated with T cell derived IFN*γ* in response to antigen stimulation^12^, suggesting that Jagged-1 may also regulate T cell responses in the human immune system. Further research is needed to determine the mechanisms responsible for the increased Foxp3 expression observed in CD4^+^ T cells in *Cd11c*^*Cre+*^*Jagged*^*ff*^*Foxp3*^*eGFP*^ animals and the consequences of this expression in immune responses.

## Materials and Methods

### Mice

*Jagged*^*ff*^ mice were the generous gift of Dr. Julian Lewis on behalf of Cancer Research UK^3^. These mice were crossed to a C57 BL/6 background 10× to ensure consistent genetic phenotype. Mice were then crossed to the *Cd11c*^*Cre-1Reiz*^ mouse from Jackson labs^29^ (stock # 008068) to make *Cd11c*^*Cre+*^*Jagged*^*ff*^ mice. In some experiments, these mice were further crossed to the *Foxp3*^*eGFP*^ mouse^30^, also purchased from Jackson labs (stock # 006769) to make a *Cd11c*^*Cre+*^*Jagged*^*ff*^*Foxp3*^*eGFP*^ strain.

All mouse work was done at the University of Michigan. Studies were approved by the University of Michigan Institutional Animal Care and Use Committee under PRO00006469, in accordance with the guidelines and regulations set forth by that regulatory body.

### BCG infection

The TICE strain of BCG was purchased from ATCC and used for these experiments. To develop an infectious inoculum, BCG was cultured at 37°C in a cell culture incubator with gentle agitation for 14 days in 7H9 broth supplemented with 10% OADC medium. Frozen stocks were prepared with 20% glycerol, 10% OADC, 50% 7H9 media and 20% of a concentrated BCG suspension. CFU of stocks were determined by plating of serial dilutions on 7H11 agar. Plates were incubated at 37°C in a cell culture incubator for 14 days prior to the enumeration of colonies. CFU of lungs was determined by homogenizing a whole mouse lung with a Tissue-Tearor in 1 mL of PBS. Lung homogenate was plated both neat and in serial dilutions and CFU enumerated as described above.

Infection of mice was done with a dose of 1.0×10^5^ CFU/mouse of BCG in 40 μL of PBS via a tongue-pull intratracheal instillation^31^. Prior to instillation, frozen BCG stocks were thawed, centrifuged at 5,000 xg for 2 minutes, and washed 3x in 1 mL of PBS before resuspending at the desired concentration.

### Cell culture

#### In vitro culture of T cells

Murine T cells were isolated using the MACS naïve T cell isolation kit II (Miltenyi) and then stimulated using 1.0×10^5^ cells/well in a 96-well flat-bottom plate. Prior to the addition of the cells, each well used for culture was incubated with 3 μg/mL anti-CD3 (Biolegend clone 145-2C11) and 3 μg/mL anti-CD28 (Biolegend clone 37.51) diluted in sterile PBS for 3 hours to coat the plate. Recombinant DLL4 (#1389-D4) and recombinant Jagged-1 (#Q63722) were purchased from R&D systems and coated on plates at the same time as anti-CD3/anti-CD28. For cultures of Tregs, 10 ng/mL of rmIL-2 and 2 ng/mL of rhTGF-β (R&D systems) were added to complete RPMI (10% FCS, 1% NEAA, 1% Na-Pyruvate, 1% Penicillin/Streptomycin, 1% L-glutamine) at the time of culture.

#### Co-culture assays

Co-culture of T cells with DCs was done using FACS sorted splenic CD11b^+^CD11c^+^ dendritic cells at a ratio of 10 naïve T cells: 1 dendritic cell. Soluble CD3 was used at a concentration of 3 μg/mL. Cultures of draining lymph node cells were performed in 96 well plates with 5.0×10^5 cells per well in 200 uL of complete RPMI. *M. bovis* PPD antigen was obtained from the USDA and used at a concentration of 10 μg/mL (Reagent code 131-B5).

#### Flow Cytometry

A single-cell suspension of lung cells was obtained by mincing whole mouse lungs in RPMI and then digesting each lung in 5 mL of RPMI + 10% FCS with 1 mg/mL Collagenase A (Roche) and 2000 U/mL of DNAse I for 45 minutes in a shaking incubator at 37°C. After incubation, tissue fragments were passed through a 5 mL syringe with an 18G needle 15-20×. Cells were centrifuged at 400 xg for 5 minutes and then washed 2x in flow cytometry buffer (500 mL PBS + 1% FCS + 2mL of 0.5 M EDTA) and resuspended in 3 mL of the same buffer. Tissue debris was removed by passing the solution filtration through 100 μM cell culture filter (Corning #352360 or similar). 100 uL of the final 3 mL suspension was used for flow cytometry analysis.

A single-cell suspension of lymph node and spleen cells was obtained by mincing the tissue and gently pushing fragments through a 100 μM filter using the plunger from a 3 mL syringe. The filter was washed extensively with flow cytometry buffer after the tissue was dispersed to remove all the cells. Cell suspensions were centrifuged and resuspended in 0.2 mL of flow cytometry buffer. 100 uL of the suspension was used for flow cytometry analysis.

Cell suspension were first stained for viability using a fixable viability stain (ThermoFisher) and then stained with antibody cocktails in 100 μL of flow cytometry buffer at room temperature with mild shaking (50-80 RPM) on an orbital shaker for 20 minutes. Dilutions of antibodies ranged from 0.2 – 2 μL/100 μL of buffer based on titration. Cells were fixed with formalin or run live through a BD LSR II flow cytometer and analyzed with FlowJo 10. Clones and dilutions of antibody were as follows: anti-CD25 clone 3C7 used at 1:200, anti-CD4 clone RM4-5 used at 1:200, anti-CD11c clone N418 used at 1:150, anti-CD11b clone M1/70 used at 1:200, anti-CD103 clone 2E7 used at 1:200, anti-Ly6C clone HK1.4 used at 1:200. All antibodies were purchased from Biolegend.

When flow-cytometry sorting was used to isolate cell populations, antibody depletion cocktails were used for pre-enrichment. Pre-enrichment for antigen presenting cells prior to sorting was done with biotinylated anti-CD3 (clone 145-2C11) and anti-CD19 (clone 6D5) both from Biolegend. Cell suspensions were incubated with antibodies for 10 minutes at room temperature at a dilution of 1:200. Each spleen was resuspended in 600 uL of flow cytometry buffer. After incubation, cells were washed 1x in 2 mL of flow cytometry buffer and then incubated with anti-biotin beads (Miltenyi) according to manufacturer instructions. Samples were then run through a MACS LS column to deplete the antibody labeled cells. Pre-enrichment for Foxp3^−^ naïve T cells was done using the MACS naïve T cell isolation kit II and then FACS sorting for GFP^−^ cells from *Foxp3*^*eGFP*^ mice.

#### Real-time PCR

RNA was isolated using TRIzol per the manufacturer’s instructions, quantified using a Nanodrop spectrophotometer, and 100 ng of RNA was reverse transcribed using iScript (Bio-Rad). Exon-spanning pre-developed assays for each of our targets were purchased from Thermo Fisher/Applied Biosystems; *Cd25* (Mm01340213_m1), *Foxp3* (Mm0047562_m1), *Tcf7* (Mm00493445_m1), *Notch2* (Mm00803077_m1). Assays were run on an Applied Biosystems TaqMan 7500 machine for 40 cycles using TaqMan master mix.

#### Measurement of cytokines in supernatants

Cytokines in cell culture supernatants were measured by Multiplex assay from Bio-Rad on a Luminex machine following the manufacturer’s instructions.

#### Statistics

All statistical analysis was performed using GraphPad Prism software.

## Author Contributions

MAS and SK designed and performed experiments and analyzed, interpreted and presented data. RA and MS performed experiments, NWL and SKL interpreted data.

## Data Availability

The datasets generated during and/or analyzed during the current study are available from the corresponding author on reasonable request.

## Acknowledgements

We would like to thank Dr. Ivan Maillard for his input and use of reagents. We would like to thank Dr. Judith Connett for editing this manuscript.

## Figure legends

**Figure S1:** A-C) Quantitation of other Notch target genes by qPCR in cells stimulated as indicated in figure 1. D-E) Viable Tregs and viable CD4^+^ T cells graphed as a function of the dose of rJagged-1.

**Figure S2:** A-B) Measurement of cytokine production from splenic CD11c^+^CD11b^+^ DC isolated by FACS from *Cd11c*^*Cre+*^*Jagged1*^*ff*^ mice compared to controls. DC were stimulated with TLR ligands for 6 hours. RNA was isolated and used for qPCR. C) Measurement of cytokine production in cell culture supernatants at 7 days after culture of naïve T cells with sorted DC from *Jagged*^*ff*^ or *Cd11c*^*Cre+*^*Jagged*^*ff*^ mice under iTreg conditions.

