## Supplementary figures and images for "Jagged-1 regulates *Foxp3* expression and cytokine production in CD4^+^ T cells"

### Supplemental figure 1

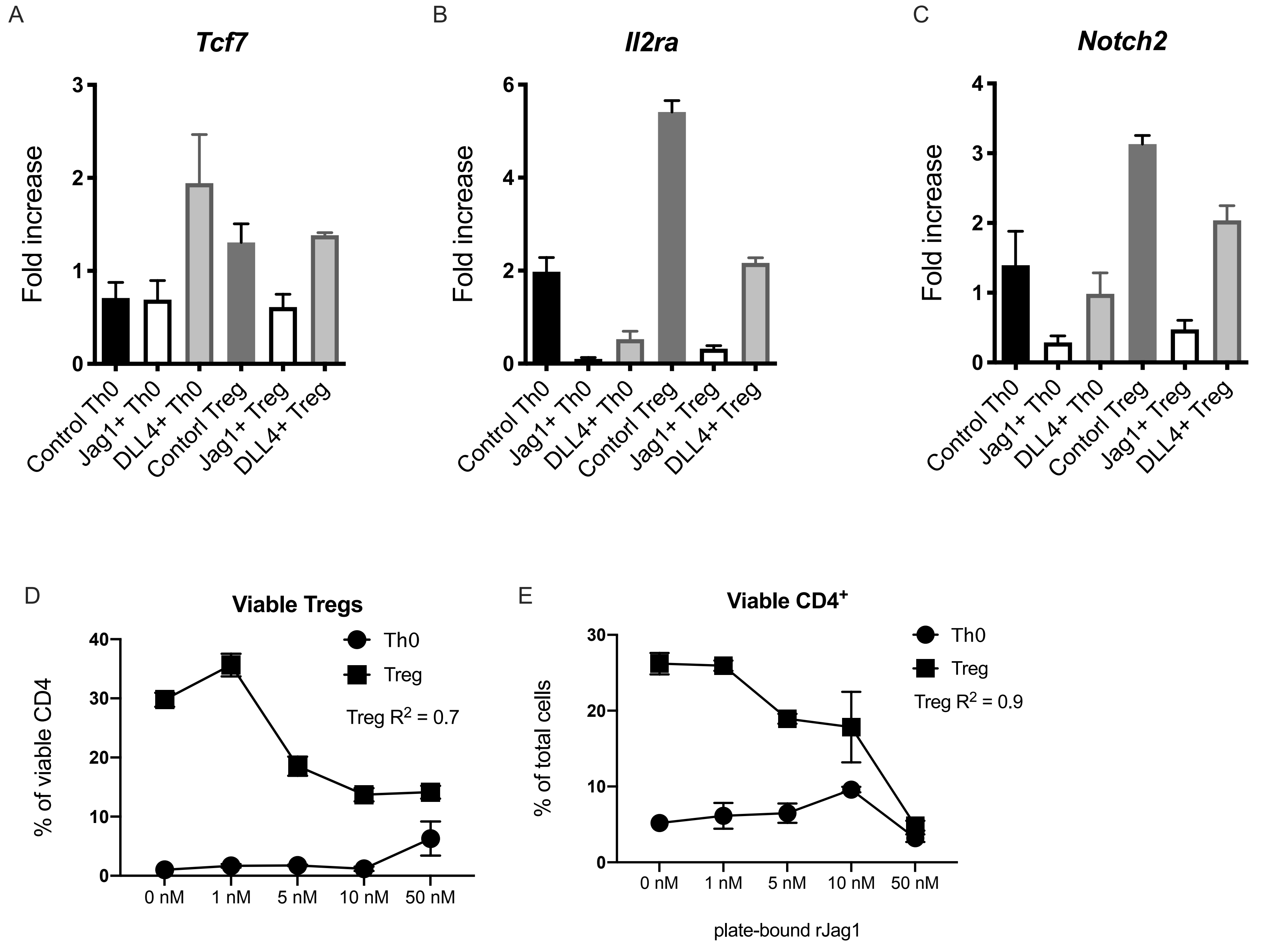

### Supplemental figure 2

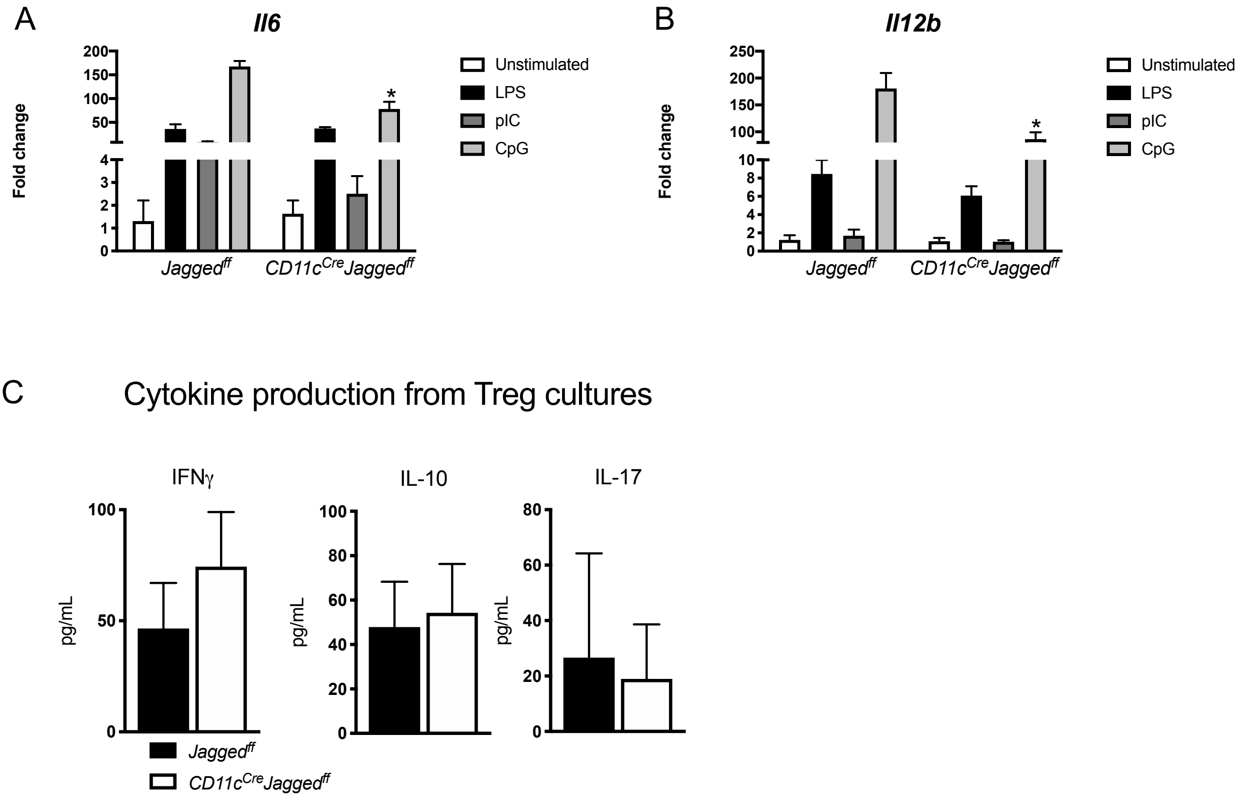
